# Quantitative comparison of the structural differences between NRAS and its mutations by well-tempered metadynamics simulations

**DOI:** 10.1101/2024.08.09.607354

**Authors:** Zheyao Hu, Jordi Marti

**Author notes:** Electronic Supplementary Information (ESI) available. See DOI: 00.0000/00000000. These authors contributed equally to this work.

## Abstract

The NRAS-mutant subset of melanoma is one of the most aggressive and lethal types associated with poor overall survival. Unfortunately, a low understanding of the NRAS-mutant dynamic behavior has lead to the lack of clinically approved therapeutic agents able to directly target NRAS oncogenes. In this work, accurate local structures of NRAS and its mutants have been fully explored through the corresponding free energy surfaces obtained by microsecond scale well-tempered metadynamics simulations. Free energy calculations are crucial to reveal the precise mechanisms of Q61 mutations at the atomic level. Considering specific atom-atom distances *d* and angles *ϕ* as appropriate reaction coordinates we have obtained free energy surfaces revealing local and global minima together with their main transitions states, unvealing the mechanisms of abnormal NRAS activation from atomic-level and quantitatively analyzing the corresponding stable states. This will help to advance in our understanding of the basic mechanisms of NRAS mutations, offering new opportunities for the design of potential inhibitors.

## 1 Introduction

RAS families play a central role in a variety of cellular processes such as signalling, survival, apoptosis or membrane trafficking^1–3^. They function like “binary switches” cycling between active (GTP-bound) and inactive (GDP-bound) states^4^. When activated, they can interact with effector proteins that control fundamental biological processes such as EGFR^5–7^, MAPK^8,9^ or PI3K ^10^. However, excessive activation of RAS will eventually lead to cancers, such as melanoma, colorectal, lung and, especially, pancreatic cancer^11,12^. Especifically, neuroblastoma RAS viral (vras) oncogene homolog (NRAS) is one of the most important RAS isoforms, with the typical mutated codon occurring at residue 61^12,13^, so that the excessive activation of NRAS is highly associated with malignant melanoma. The NRAS Q61-mutant subset of melanoma shows aggressive clinical behaviour, poor prognosis and low overall survival annually causing the largest number of deaths^14–17^.

It has been observed that NRAS mutations tends to stay at its active GTP-bound state as its most predominant confiormation^18^, mainly due to the mutations are codon 61 impairing the intrinsic enzymatic activity of NRAS to hydrolyze GTP to GDP. This makes the task of drugging NRAS very difficult. Although several studies ^19–24^ have been focused on the most relevant NRAS mutations and some immunotherapies are available to patients with cancers related to NRAS oncogenes^15^, there is still important lack of knowledge on the detailed atomic level structure of NRAS and, especially, of efficient strategies capable of directly targeting NRAS^25–27^. For instance, some strategies such as the potential inhibitor HM-387 have been recently designed *in silico*, in order to target the NRAS-Q61 positively charged mutant R61, inducing the GTP-bound NRAS-Q61 oncogenic mutations to an “off-like” state ^24^. However, its effectiveness still needs to be verified in wet lab and in clinical practice. Therefore, the detailed conformation and local structure of NRAS still requires significant increase of knolwledge, in order to be used for locating potential targetting pockets that may facilitate the discovery and improvement of potential drug-like compounds.

Detailed information on atomic interactions and local structures at the all-atom level of NRAS are of crucial importance in inhibitor development^28^, given the difficulty to access nanoscale length and time in experimental measurements. In a previous work, the impact of NRAS-Q61 mutations on its structural characteristics was qualitatively investigated ^24^ by means of microsecond time-scale molecular dynamics simulations. In this regard, quantitative properties such as free energy barriers associated with Q61 mutations and multidimensional free energy (hyper)surfaces (FES) of NRAS and its mutations in solution are still remain to be revealed. Moreover, the choice of appropriate collective variables (CV) to accurately describe the dynamic behavior of NRAS remains unclear. To move forward in this direction, we report in the present work two high-quality CV, which will be one cornerstone of the present and future RAS investigations ^29–31^. Overall, combined with well-tempered metadynamics (WTM) simulations, we report in the present work a series of FESs obtained in aqueous solution for the GTP-bound wild-type NRAS and its mutations, helping to foster subsequent medicinal development in wet lab experiments and in clinical tests.

## 2 Methods

### 2.1 System preparation and setups

In the present work we conducted WTM simulations of 5 NRAS isoforms with the same sequences represented in the previous work^24^. Each system contained one isoform of the GTP or GDP bound NRAS complex fully solvated by TIP3P water molecules^32^ in potassium chloride (0.15 M) and magnesium chloride (0.03 M) solution. The detailed number of particles in the simulation boxes are listed in Table 1. All WTM inputs were generated by means of the CHARMM-GUI solution builder^33–35^ assuming the CHARMM36m force field^36^. All bonds involving hydrogens were set to fixed lengths, allowing fluctuations of bond distances and angles for the remaining atoms. Crystal structures of GDP-bound NRAS and GTP-bound NRAS were downloaded from RCSB PDB Protein Data Bank^37^, namely file names “6zio” and “5uhv”. The sets of NRAS proteins (wild type, Q61R, Q61K and Q61L) were solvated in a water box, all systems were energy minimised and well equilibrated (NVT ensemble) before generating the WTM simulations.

**Table 1.**
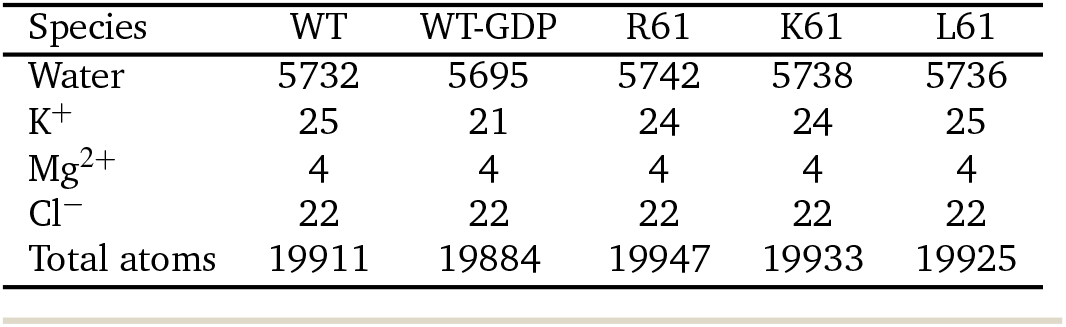
Number of particles in the GTP-bound NRAS simulation boxes.

| Species | WT | WT-GDP | R61 | K61 | L61 |
| --- | --- | --- | --- | --- | --- |
| Water | 5732 | 5695 | 5742 | 5738 | 5736 |
| K<sup>+</sup> | 25 | 21 | 24 | 24 | 25 |
| Mg<sup>2+</sup> | 4 | 4 | 4 | 4 | 4 |
| Cl<sup>-</sup> | 22 | 22 | 22 | 22 | 22 |
| Total atoms | 19911 | 19884 | 19947 | 19933 | 19925 |

The GROMACS/2021 package (version released on January 28th, 2021) was employed^38^ for the system minimization and equilibration steps. The system was minimised with steepest descents and a conjugate-gradient step every 10 steps until convergence to Fmax < 1000 kJ/mol. After energy minimisation, a time steps of 1 fs was used in all equilibration simulations and the particle mesh Ewald method with Coulomb radius of 1.2 nm was employed to compute long-ranged electrostatic interactions. The cutoff for Lennard-Jones interactions was set to 1.2 nm. The system was equilibrated for 1.25 *µ*s with the Nosé-Hoover thermostat at 310.15 K in the NVT ensemble. Periodic boundary conditions in three directions of space have been taken.

### 2.2 Well-tempered metadynamics

A wide variety of methods has been proposed to handle the complex problem of computing free energy landscapes in multidimensional systems^39–43^. In this work we have employed WTM, a method able to efficiently explore free energy surfaces of complex systems using multiple reaction coordinates, very successful for a wide variety of complex systems^29,44–48^. The main advantage of WTM is its suitability for a wide variety of systems, including model cell membrane systems with attached small-molecules and proteins^49–52^, recently developed in our group.

After equilibration we switched to run another 3.0 *µ*s of WTM simulations for each system to perform Gibbs free energy calculations, starting from the last configuration of MD simulations with the Nosé-Hoover thermostat at 310.15 K and the Parrinello-Rahman barostat at 1 atm in the NPT ensemble. Long well-tempered metadynamics simulations were performed using the joint GROMACS/2022.5-plumed-2.9.0 tool^53,54^. Periodic boundary conditions in the three directions of space were also considered. The set of CVs adopted in the WTM simulations will be discussed in full details below (see Section 3.1).We should point out that determining and using more than two CVs in a WTM simulation is a hard challenge from the computational point of view and would produce a four-dimensional FES, unsuitable to deal with. A very usual choice is to consider one or two CVs^55^. From our experience, the use of two CVs produces a complete enough description of the FES and it is the optimal way to proceed. Later, it is a usual procedure to project the free energies onto one single coordinate, integrating out the contribution of the second CV. The values for the parameters of the metadynamics simulations^56^ are listed in Table 2.

**Table 2.**
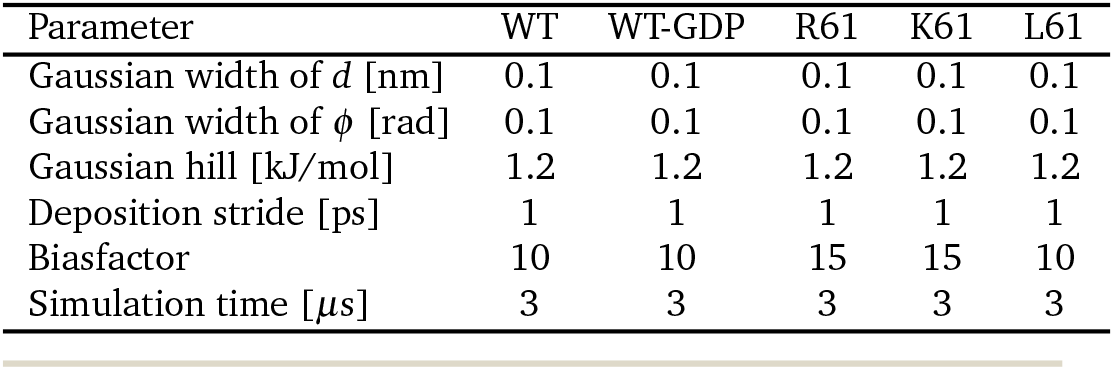
WTM simulation parameters of GTP-bound NRAS systems.

| Parameter | WT | WT-GDP | R61 | K61 | L61 |
| --- | --- | --- | --- | --- | --- |
| Gaussian width of $d$ [nm] | 0.1 | 0.1 | 0.1 | 0.1 | 0.1 |
| Gaussian width of $\phi$ [rad] | 0.1 | 0.1 | 0.1 | 0.1 | 0.1 |
| Gaussian hill [kJ/mol] | 1.2 | 1.2 | 1.2 | 1.2 | 1.2 |
| Deposition stride [ps] | 1 | 1 | 1 | 1 | 1 |
| Biasfactor | 10 | 10 | 15 | 15 | 10 |
| Simulation time [μs] | 3 | 3 | 3 | 3 | 3 |

### 2.3 Data analysis and visualisation

The R-package metadynminer was used to analyze the WTM results, such as calculating the free energy surface, finding minima and analyzing transition pathways between stable states (minimum free energy paths) in order to locate the transition states of the system^57^. The software VMD^58^ was used for visualisation purposes.

## 3 Results

In a previous work^24^, the effect of Q61 mutations on the microscopic configurations of NRAS was revealed at the atomic level. In the case of GTP-bound NRAS, we found that the positively charged mutations Q61R and Q61K maintain stable conformations comparable to wild-type NRAS and that the mutated residues R61 and K61 showed significant interaction properties with guanosine triphosphate (GTP) whereas, in a full qualitatively different fashion, the mutation NRAS-Q61L (with the non-polar residue L61) can be captured by a hydrophobic pocket. Limited by unbiased simulations, the free energy barriers of these interactions cannot be accurately measured. Besides, the FES of NRAS have not been reported yet. This information would be of crucial importance in order to fully understand the signaling of NRAS and the impact of Q61 mutations on it, as well as for subsequent pharmaceutical research and development. In order to access clear and well defined FES, it is of paramount importance to know the best reaction coordinates or CV, to be employed in precise calculations. In the present work, we will evaluate the best CV candidates for NRAS and its mutations in aqueous ionic solution, and evaluate their reliability by computing the FES in each case. Moreover, the physical meaning of the results will be discussed and the proof of convergence of the WTM simulations will be reported in the Supporting Information (SI).

### 3.1 Well-tempered metadynamics simulations: collective variables

The enhanced sampling method WTM applies a time-dependent biasing potential along a set of CV by adding a Gaussian additional bias potential to the total potential in order to overcome barriers larger than *k*_*B*_*T*, with *k*_*B*_ being Boltzmann’s constant and *T* the temperature. In the method, sampling is performed on a selected number of degrees of freedom (*s*_*i*_), i.e. a chosen set of CV. For each degree of freedom, the biased potential *V* (*s*_*i*_, *t*) is a dynamical function constructed as the sum of Gaussian functions^56,59,60^:

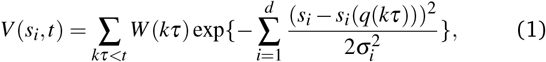

where *k* is an integer, *τ* is the Gaussian deposition stride, *W* (*kτ*) is the height of the Gaussian and *σ*_*i*_ is the width of the Gaussian for the i-th CV. Unlike unbiased molecular dynamics simulations, able to track the dynamical evolution of the system around equilibrium states (stable ststes), the biased potential can force the system to move around all possible states inside a particular range of the subspace of selected CV. In the present work, we have taken a specific approach similar to previous works^49,52,61^ where distances combined with angles were chosen (see Fig. 1).

**Fig. 1.**
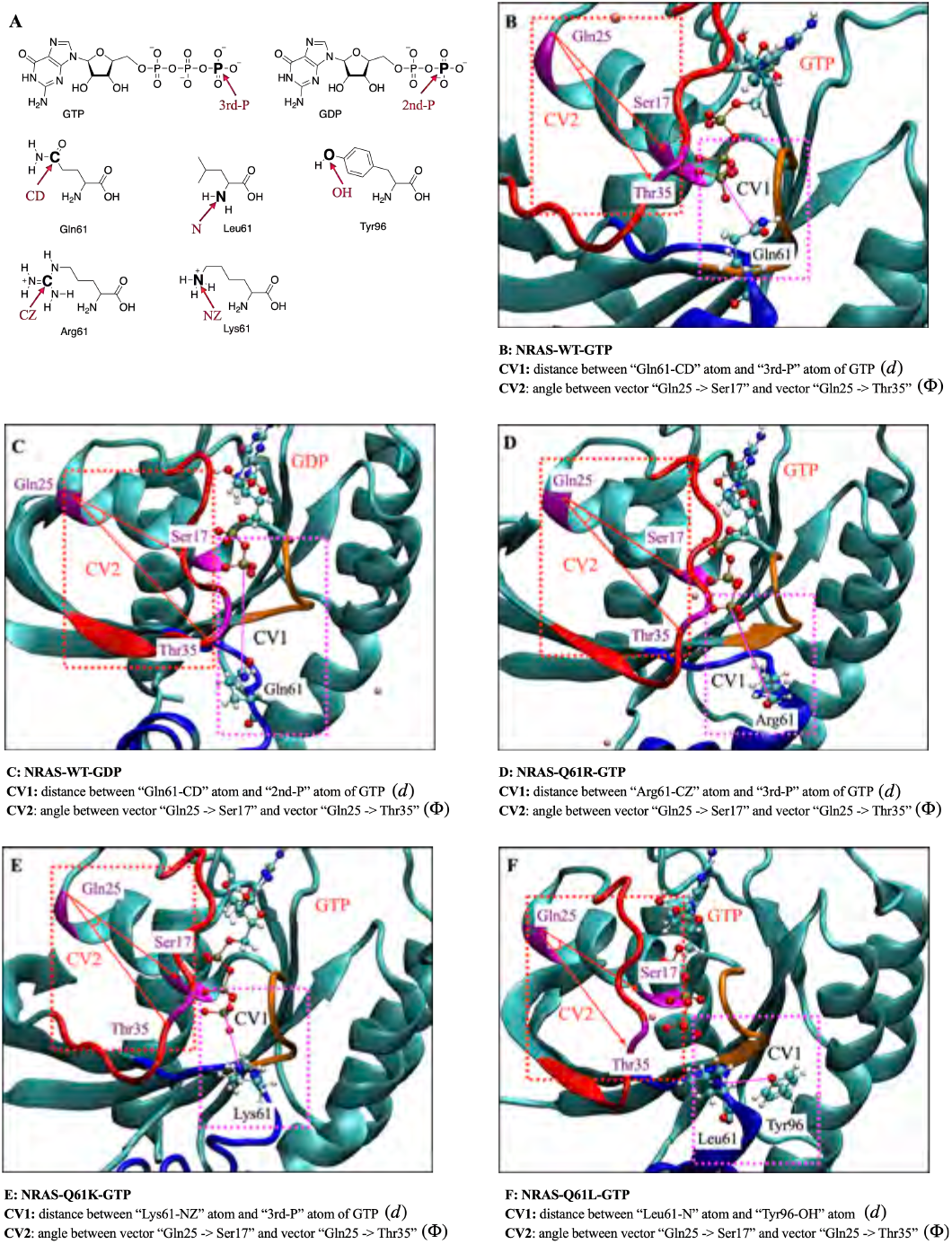
CV sets employed in the present work: (A) Schematic diagram of the atoms involved in the definition of CVs; (B) The CVs used for GTP-bound wild-type NRAS are: (1) the distance (*d*) between “Gln61-CD” atom and “3rd-P” atom of GTP as CV1 and the angle (*ϕ*) between vector “GIn25-C*α* -> Ser17-C*α*” and vector “GIn25-C*α* -> Thr35-C*α*” as CV2; (C) for GDP-bound wild-type NRAS; (D) for GTP-bound NRAS-Q61R; (E) for GTP-bound NRAS-Q61K; (F) for GTP-bound NRAS-Q61L, the distance between “Leu61-N” atom and “Tyr96-OH” atom is taken as CV1 (*d*) whereas CV2 is same as in the previous cases.

The conformational dynamics and biological effects of RAS families are mainly reflected by the structures of Switch-I (SWI, residues 27-37) and Switch-II (SW-II, residues 58-64)^24,62–64^. So the CVs chosen in this work are directly related to SW-I and SW-II will help to fully describe all intermediate states of NRAS and its mutants.

After a thorough selection from a variety of distances and angular coordinates, the two CV selected to perform the 2D WTM calculations are (see Fig. 1): (1) distances (CV1 *d*) between the 61st amino acid residue of NRAS and the terminal phosphate group of GTP/GDP. In the case of NRAS-Q61L, considering that the non-polar hydrophobic amino acid Leu61 cannot form stable interaction with the negatively charged phosphate group, we instead used the distance between Leu61 and Tyr96 as CV1; (2) the angles between two vectors (CV2 *ϕ*_*i*_) formed by the vector “GIn25-C*α* -> Ser17-C*α*” and vector “GIn25-C*α* -> Thr35-C*α*” in all cases. Among them, CV1 (*d*) characterizes the biological function of Q61 and the impact of Q61 mutations on the intrinsic GTPase function of NRAS and CV2 (*ϕ*) represent the “on/off” status of switch-I well. The fully detailed information of each CV is described in Fig. 1. The averaged results for each CV are reported in the following Sections 3.2 and 3.4.

### 3.2 The two-dimensional free energy landscapes of NRAS wild-type and its main Q61 mutations

Two dimensional (2D) free energy landscapes of NRAS and its mutations are represented in Fig. 2 for the two selected CV. We can observe that several main basins (minimum, such as A, B …) and their corresponding molecular conformations are well defined in all cases. We have only indicated the basins which are clearly seen from the FES, assuming that even with well converged runs the clearest definition of the FES would require a huge amount of statistics of the order of 20 *µ*s, for each WTM simulation, out of the standard available ranges. The coordinates of the representative basins corresponding to the Fig. 2 are reported in Table 3, whereas the detailed coordinates of minima are reported in SI-Table 1 to SI-Table 5.

**Table 3.**
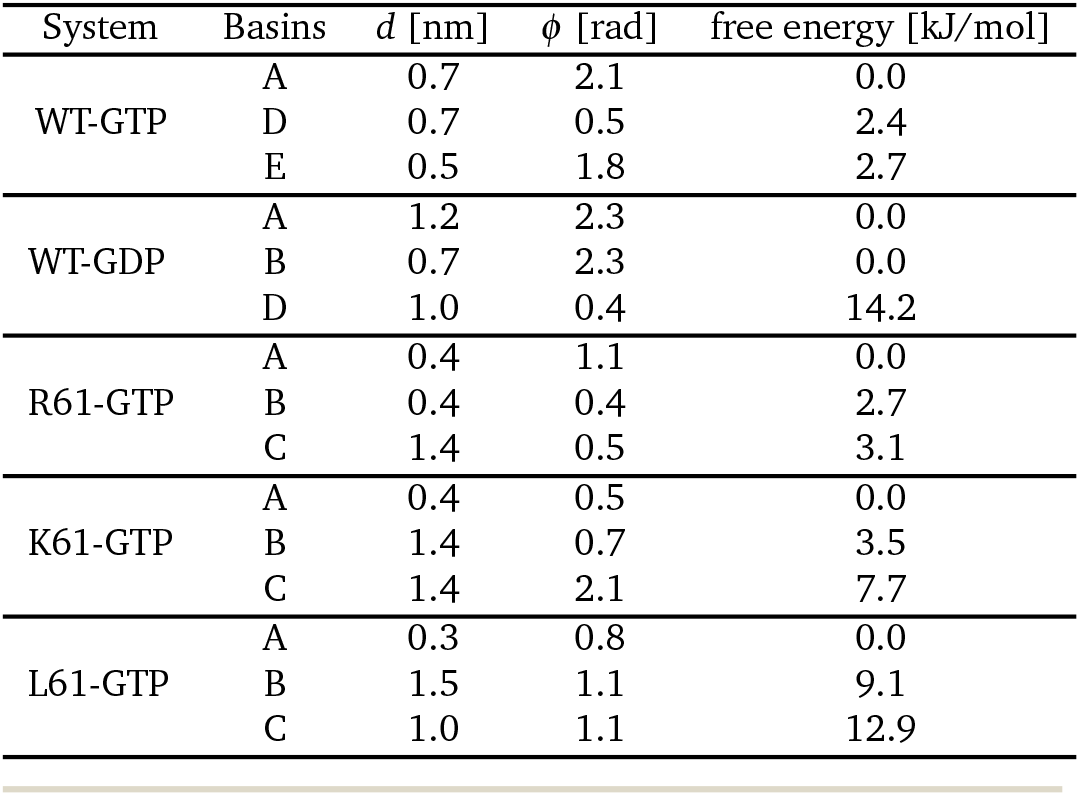
The coordinates of the representative basins (minima) corresponding to the Fig. 2.

| System | Basins | $d$ [nm] | $\phi$ [rad] | free energy [kJ/mol] |
| --- | --- | --- | --- | --- |
| WT-GTP | A | 0.7 | 2.1 | 0.0 |
|  | D | 0.7 | 0.5 | 2.4 |
|  | E | 0.5 | 1.8 | 2.7 |
| WT-GDP | A | 1.2 | 2.3 | 0.0 |
|  | B | 0.7 | 2.3 | 0.0 |
|  | D | 1.0 | 0.4 | 14.2 |
| R61-GTP | A | 0.4 | 1.1 | 0.0 |
|  | B | 0.4 | 0.4 | 2.7 |
|  | C | 1.4 | 0.5 | 3.1 |
| K61-GTP | A | 0.4 | 0.5 | 0.0 |
|  | B | 1.4 | 0.7 | 3.5 |
|  | C | 1.4 | 2.1 | 7.7 |
| L61-GTP | A | 0.3 | 0.8 | 0.0 |
|  | B | 1.5 | 1.1 | 9.1 |
|  | C | 1.0 | 1.1 | 12.9 |

**Fig. 2.**
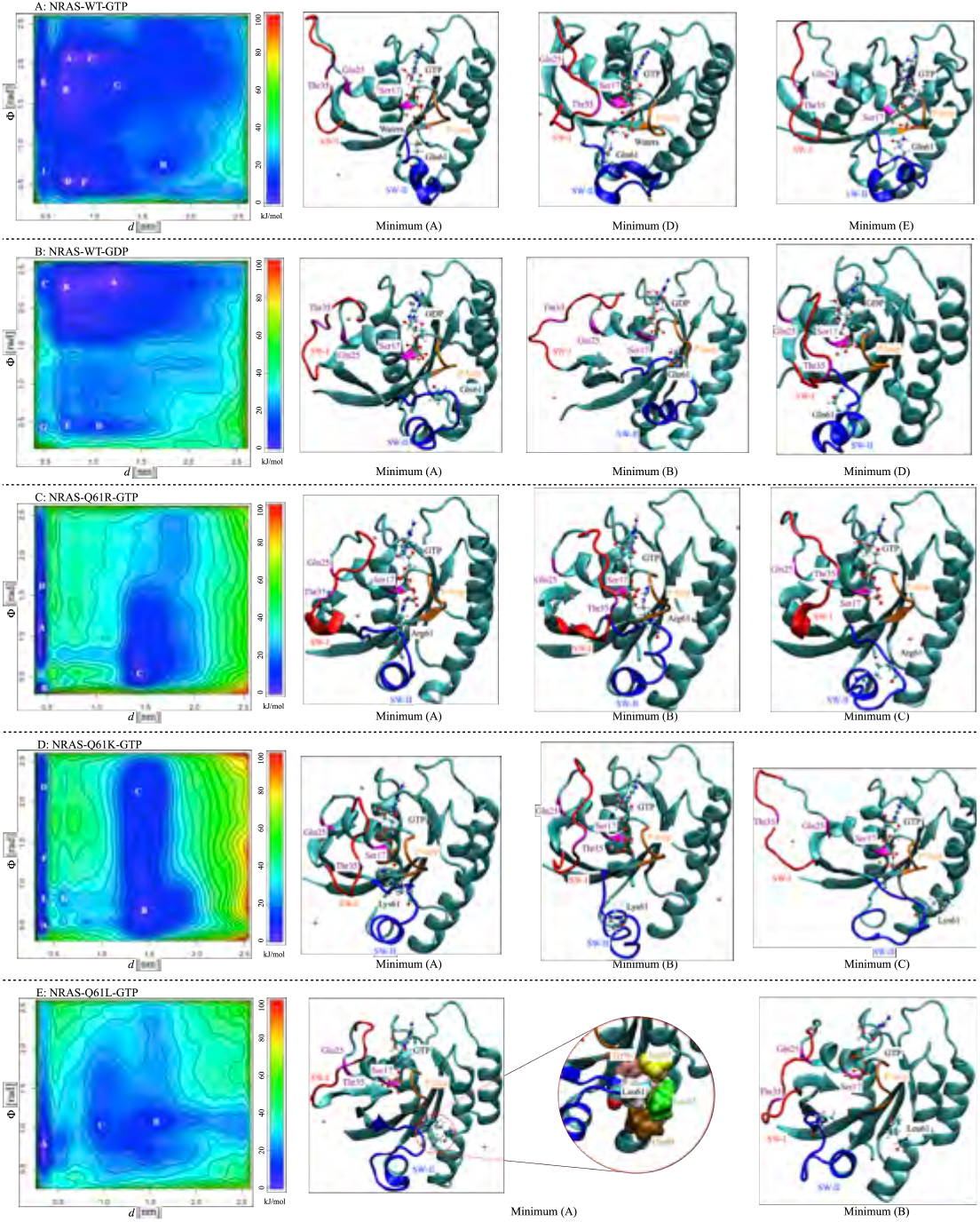
The 2D free energy landscape F(*d, ϕ*) (kJ/mol) for the wild-type NRAS and its Q61 mutations. Basins “A”, “B”, “C”, “D”, “E”, “F”, “G”, “H” and “I” are the stable states (minima) and are depicted in white. The conformations corresponding to the representative basins are listed on the right side of the 2D free energy landscape. The coordinates of representative basins (minima) of the 2D free energy landscape are listed in Table 3, and the detailed coordinates of minima are reported in SI-Table 1 to SI-Table 5.

As a general concept, the basins in the FES indicate the most probable configurations of the system in equilibrium. As a matter of fact, WTM is able to reveal those stable configurations in such a way that the free energy barriers that the system needs to overcome to shift between stable states can be estimated with a high degree of accuracy. In this work, we can distinguish several basins for each simulation system proposed. In the case of GTP-bound wild-type NRAS, several stable states can be clearly observed (see Fig. 2A). GTP-bound wild-type NRAS is responsible for the normal biological signal transduction. The corresponding FES indicates that the conformation of wild-type NRAS is evenly distributed and that free energy barriers required to switch between several stable states are low magnitude. The most stable conformation is distributed between minima A, B and C. The representative conformation minimum A holds the SW-I half open, while SW-II remains compact. The residue Q61 stabilizes the water molecules around *γ*-phosphate through hydrogen-bond interactions (HB) and it is therefore responsible for the intrinsic GTP hydrolysis activity of NRAS. In the meta-stable state D, the SW-I is closed and the SW-II (especially residue Q61) holds a similar fashion with its configuration in minimum A. The above results indicate that Q61 can stabilize water molecules around *γ*-phosphate of GTP through HB. This is fully consistent with previously reported mechanisms indicating that residue Q61 participates in the intrinsic GTP hydrolysis activity of NRAS by stabilizing transient hydronium ions around the *γ*-phosphate^19–24^, although unbiased classical MD simulations rather than QM/MM simulations were used in those investigations.

From our results we observe that in the GDP-bound wild-type NRAS case, both SW-I and SW-II are kept open, with a free energy difference between basins A, B and D of approximately 14.2 kJ/mol, as shown on Fig. 2B and Table 3. In general, the release of the *γ*-phosphate after GTP hydrolysis allows the two switch regions to relax into a more “open” state, which is in good agreement with the so called “GDP-specific conformation”^4,62,65,66^.

Mutations in NRAS Q61 significantly reduce the intrinsic GTPase activity, leading to abnormal activation and ultimately causing cancer^67–70^. In a previous work^24^, we revealed the effect of Q61 mutations on the conformational characteristic of GTP/GDP-bound NRAS, while the global FES and the precise free energy barriers were estimated^24^. In the GTP-bound NRAS Q61 positively charged mutations (Q61R and Q61K), the FES shows significant differences compared to GTP-bound wild-type NRAS, Fig. 2 C and D. The positively charged Q61 mutations show a marked tendency to maintain the SW-I/II regions in a compact conformation. In the GTP-bound NRAS-Q61R case, the basins A, B at the FES correspond to same distance *d* (0.4 nm) and two different angles (1.1, 0.4 rad), being this in good agreement with the previous results obtained from unbiased MD simulations^24^.

Furthermore, large free energy barriers of the system are explored by WTM so that additional undiscovered meta-stable state can be found. For instance, the minimum C at the FES corresponds to the distance *d* = 1.4 nm and angle *ϕ* = 0.5 rad. In the case of the GTP-bound NRAS-Q61K, due to the similar chemical structures and charge properties between R61 and K61, their FES are quite similar. The basin A at the FES corresponds to the distance *d* = 0.4 nm and angle = 0.5 rad, whereas the meta-stable state B corresponds to the distance *d* = 1.4 nm and angle *ϕ* = 0.7 rad. In addition, since the side chain length of K61 is slightly smaller than that of R61, the interaction strength between K61 and GTP is slightly smaller than that of R61, making the largely opened conformation of NRAS-Q61K easier to reach in comparison with NRAS-Q61R, providing a stable basin C with the coordinates (1.4 nm, 2.1 rad). Finally, for the non-polar mutation Q61L, hydrophobic interactions of L61 with the hydrophobic pocket composed by the D92, L95, Y96 and Q99^24^ can be also detected in the FES stable basin A with the coordinates (0.3 nm, 0.8 rad) (see Fig. 2 E). Moreover, benefiting from the WTM, an unknown meta-stable state B (1.5 nm, 1.1 rad) corresponding to the dissociation of L61 from the hydrophobic pocket has been also identified.

### 3.3 The conformational transitions of GTP-bound NRAS wild-type, NRAS-Q61R and NRAS-Q61L

The barriers between stable state basins indicate the amount of free energy required for NRAS to exchange its conformation between stable configurations. Using the metadynminer method^57^, we have been able to trace the minimum free energy path (MFEP) and calculate the energy barriers ΔF with high accuracy. Metadynminer tracks the MFEP by iteratively refining the path connecting stable states, which converges to the minimum free energy trajectory between them. Furthermore, the transition states (TS) between two selected stable states can also be determined from the knowledge of MFEP. To characterize the conformational change differences in FES reported in Fig. 2, we have used the MFEP to explore the transition between different conformation changes of NRAS and its mutants. By means of comparing the effects of Q61 mutations on the local structures of the well-established potential binding pockets (SW-I/II), we may provide initial global structural differences that might help design anti-cancer drugs targeting the (meta)stable states reported in this work. The MFEP of GTP-bound NRAS and the corresponding main TS are reported in Fig. 3.

**Fig. 3.**
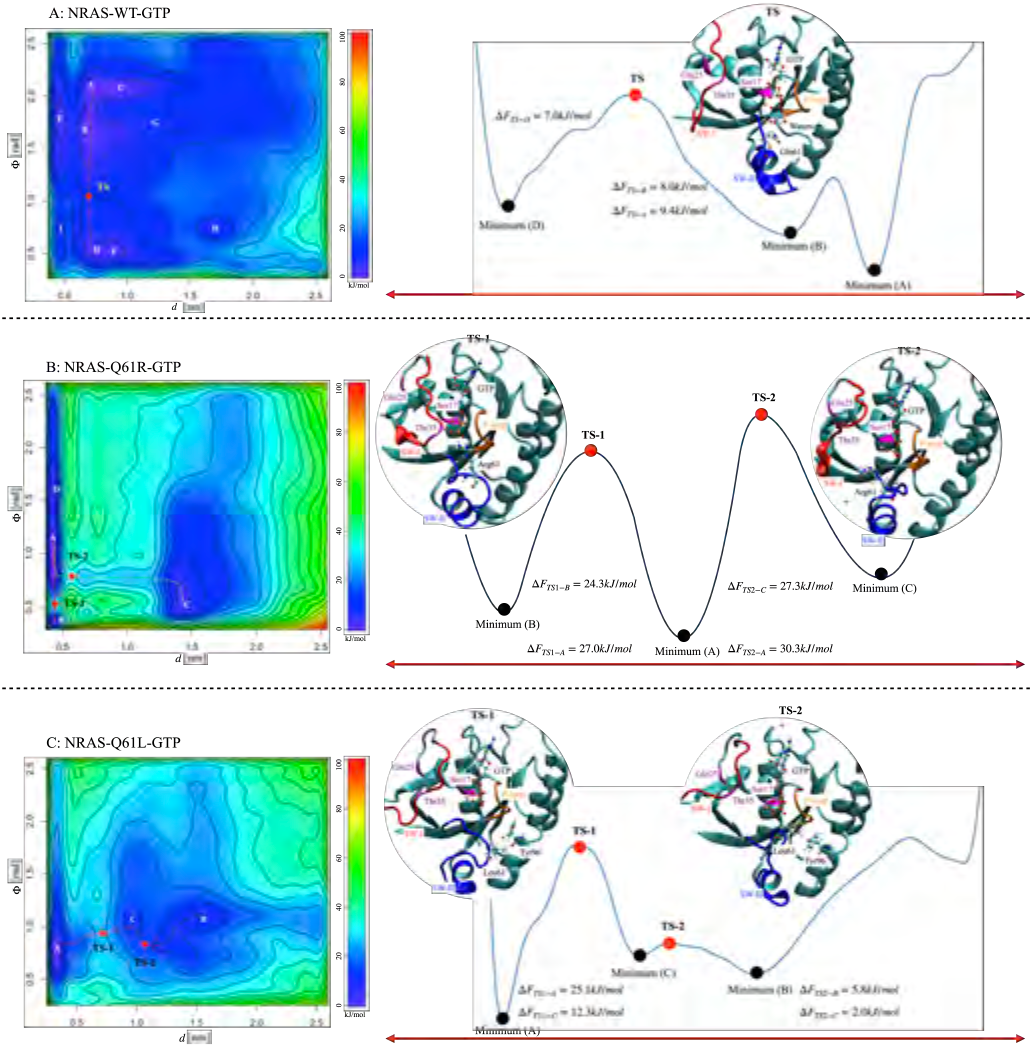
The minima free energy path (MFEP) between selected stable states. (A) The MFEP of GTP-bound wild-type NRAS. Left, the MFEP between the selected minima A/D; right, the schematic diagram of MFEP, free energy barrier ΔF and the corresponding conformation screenshot of the TS. (B) The MFEP of GTP-bound NRAS-Q61R. Left, the MFEP between the selected minima A/B (depicted in magenta) and minima A/C (depicted in yellow); right, the schematic diagram of MFEP, free energy barrier ΔF and the corresponding conformation screenshot of the TS. (C) The MFEP of GTP-bound NRAS-Q61L. Left, the MFEP between the selected minima A/C and minima C/B; right, the schematic diagram of MFEP, free energy barrier ΔF and the corresponding conformation screenshot of the TS.

The FES minima of GTP-bound wild-type NRAS was discussed in the above Section 3.2. The two large free energy basins are mainly composed of minima A, B, C and minima D, F respectively. Consequently, minima A and D have been selected to represent the two stable states of GTP-bound wild-type NRAS and employed to calculate the MFEP between them (see Fig. 3 A). The free energy differences between basins A and D has been estimated to be of 2.4 kJ/mol. Along the MFEP depicted in Fig. 3 A, we identified free energy barriers ΔF_*TS*−*A*_ from “A” to “TS” of 9.4 kJ/mol, ΔF_*TS*−*B*_ from “B” to “TS” of 8.0 kJ/mol and ΔF_*TS*−*D*_ from “D” to “TS” of 7.0 kJ/mol. From the MFEP and the screenshots of TS, the conformational changes of GTP-bound wild-type NRAS are mainly reflected in the “on/off” states of SW-I, while the Q61 continues to stabilize the water molecules around *γ*-phosphate, again in good agreement with the hydrolysis mechanism of GTP^19–24^.

Considering the similarities between the NRAS positively charged mutations (R61 and K61), GTP-bound NRAS-Q61R will be taken as an example to introduce the effect of positively charged Q61 mutations on the conformational transitions. In GTP-bound NRAS-Q61R case, at least three stable states (minima A, B and C) can be clearly detected. The free energy difference between basins B and C has been estimated to be of 0.4 kJ/mol (see Table3), so two MFEPs have been calculated to characterize the GTP-bound NRAS-Q61R conformational changes, namely MFEP_*A*−*B*_ and MFEP_*A*−*C*_ (see Fig. 3B). Along the MFEP_*A*−*B*_, the free energy barrier ΔF_*TS*1−*A*_ from “A” to “TS1” is of 27.0 kJ/mol and ΔF_*TS*1−*B*_ from “B” to “TS1” is of 24.3 kJ/mol, so that the conformational TS1 is mainly correlated to the “on/off” state of SW-Correspondingly, along the MFEP_*A*−*C*_, the free energy barrier ΔF_*TS*2−*A*_ from “A” to “TS2” is of 30.3 kJ/mol and ΔF_*TS*2−*C*_ from “C” to “TS2” is of 27.3 kJ/mol, so that the conformational TS2 is related to both SW-I and SW-II. Moreover, due to the significantly high TS free energy barriers ΔF, NRAS is “locked” in a small conformational region.

Conversely, the GTP-bound NRAS-Q61L case is much simpler than the previous case, with three stable states (minima A, B and C) to be considered for the calculation of the MFEP (see Fig. 3C). Basin A is the most stable state, with free energy differences with basins B and C of 9.1 kJ/mol and 12.9 kJ/mol respectively (see Table3), significantly larger than in the previous R61 case. Along the MFEP_*A*−*C*_, the free energy barriers are: ΔF_*TS*1−*A*_ from “A” to “TS1” of 25.1 kJ/mol and ΔF_*TS*1−*C*_ from “C” to “TS1” of 12.3 kJ/mol. Moreover, along the MFEP_*B*−*C*_, the free energy barriers are moderately smaller: ΔF_*TS*2−*B*_ from “B” to “TS2” is of 5.8 kJ/mol and ΔF_*TS*2−*C*_ from “C” to “TS2” is of 2.0 kJ/mol. The physical meaning of MFEP between minima A, C and B relies on the binding and dissociation states of L61 with the hydrophobic pocket.

### 3.4 One-dimensional free energy profiles

From the 2D free energy landscapes and minimum free energy paths obtained from WTM simulations (Figs. 2 and 3) it is possible to obtain a one-dimensional (1D) free energy profile where one single CV is considered and the second one has been integrated out. This calculation along one single CV (*s*_*i*_, *i* = 1, 2), allows us to directly compare free energy barriers to experimental findings, as it was pointed out by Jämbeck et al.^71^. These authors pointed out that a direct route to connect FES to experiments is normally cumbersome, since information about binding modes in solutes obtained from experiments is normally not available. Then, it is possible to use standard binding free energies Δ*G*_*bind*_ determined from experiments as an indirect measurement of the accuracy of computed free energy barriers. In the present work, 1D free energy profiles F(*s*_1_) (equivalent to the Δ*G*_*bind*_ mentioned above) can be obtained as^71,72^:

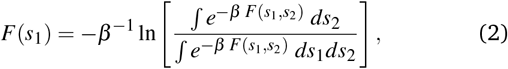

where *s*_1_ and *s*_2_ are the CV, *β* = 1*/*(*k*_B_*T*), *k*_B_ is the Boltzmann constant and *T* is the absolute temperature. This means that all possible paths for the CV labelled as *s*_2_ have been integrated out and averaged. In the present case, the results of Fig. 4 reveal a series of free energy barriers around the basins. We have obtained clear asymmetry for all free energies dependent of *d* and *ϕ*, indicating the existence of distinguishable distributions for both distances and angles.

**Fig. 4.**
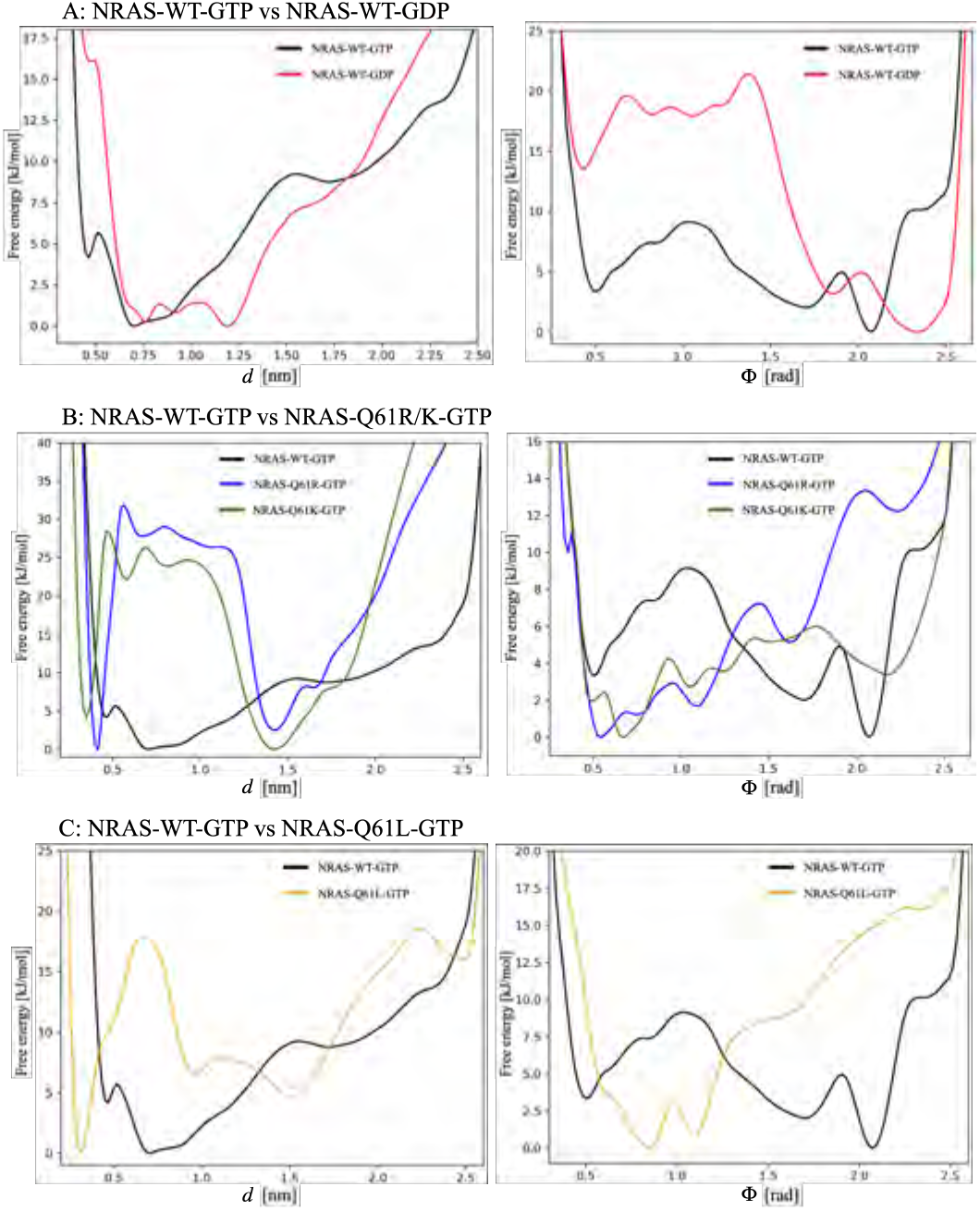
1D free energy profiles for two CV (*d, ϕ*). Top row correspond to: (A) The comparison of 1D free energy profiles between GTP-bound wild-type NRAS and GDP-bound wild-type NRAS; The second row correspond to: (B) The comparison of 1D free energy profiles between GTP-bound wild-type NRAS and GTP-bound NRAS-Q61R/K; and the last row correspond to: (C) The comparison of 1D free energy profiles between GTP-bound wild-type NRAS and GTP-bound NRAS-Q61L. In each row, the left column corresponds to the distances CV *d*, whereas the right column corresponds to the angles CV *ϕ* defined in Figure 1 for each NRAS. In order to directly compute the height of free energy barriers, each absolute minimum has been set equal to zero.

As a general fact, both angles *ϕ* and distances *d* characterize well the movement of the corresponding NRAS functional regions SW-I/II. The barriers for the free energies dependent of angles *ϕ* and distances *d* are in the range of 9 kJ/mol the largest in the GTP-bound wild-type NRAS case, and the free energies of distances *d* minimum tend to be located at the coordinate 0.7 nm, where Q61 plays the role of stabilizing the water molecules around *γ*-phosphate. When GTP is hydrolyzed to GDP, the coordinates corresponding to the free energy minima of *ϕ* and *d* both increased to 2.3 rad and 1.2 nm respectively (see Fig. 4A). The barriers for the free energies dependent of *ϕ* are in the range of 22 kJ/mol the largest.

The GTP-bound NRAS-Q61 mutations show the opposite trend in comparison with the wild-type NRAS, as reported in Fig. 4B and C. The barriers for the free energies dependent of angles *ϕ* are similar with those obtained from wild-type NRAS, say around 6 to 13 kJ/mol, while the free energy minima coordinates are located in the smaller position: 0.5 rad for NRAS-Q61R, 0.7 rad for NRAS-Q61K and 0.8 rad for NRAS-Q61L. As for the distances *d*, there exist higher free energy barriers that force the system to remain at the smaller coordinate position. The free energy barriers for the R61/K61 interaction with *γ*-phosphate are 32 kJ/mol and 25 kJ/mol, respectively, and the free energy barrier for the L61 interaction with the hydrophobic pocket on the *α*-helix 3 is 18 kJ/mol. These numbers clearly indicate that after mutation, the NRAS conformational changes should be attributed more to fluctuations of the distances *d* than to changes in coordinate angles *ϕ*.

## 4 Conclusions

NRAS are small GTPases with the ability to regulate cell growth, differentiation and survival. The mutated NRAS shows the strong association with the malignant melanoma, which has an aggressive clinical behavior and poor prognosis, as well as lower overall survival. Several efforts have been focused on the structural and pharmaceutical aspects of NRAS-Q61 mutations, but validated inhibitors are still missing. There is also a lack of accurate FESs of NRAS and its mutants. The present study is an extension of previous investigations^24^, where the WTM has been employed to evaluate the FES of NRAS and its Q61 mutations in detail. Their conformational differences in solution will help us understand the impact of the NRAS Q61 mutations and further promotes the inhibitor design. In this work, we have conducted WTM simulations of five NRAS systems and obtained the corresponding FES, considering in all cases a specific pair of reaction coordinates or collective variables. Our simulations reached the scale of 3.0 *µ*s, for a total of 15.0 *µ*s. In all cases the convergence of the WTM simulations has been clearly proved. With the help of well-tempered metadynamics simulations we have calculated 2D FES and the 1D profiles after integrating out one CV. All conformationally accessible regions of NRAS and its mutations have been fully explored. We observed that each system has several stable states, which are the signature of the equilibrium configurations for each system. The two selected CV are: (1) a distance *d* between the 61st aminoacid residue of NRAS and the terminal phosphate group of GTP/GDP and (2) the angle *ϕ* between the two vectors defined by “GIn25-C*α* -> Ser17-C*α*” and “GIn25-C*α* -> Thr35-C*α*”.

In the 2D FES of GTP-bound wild-type NRAS, a series of minima (“A” to “I”) were detected with conformations evenly distributed within the low free energy barriers. The main feature of the conformations corresponding to minima A, B and D is the existence of “bridge water”, which are stabilized between Q61 and *γ*-phosphate by hydrogen bonding interactions with them. This is consistent well with the reported intrinsic GTP hydrolysis mechanism of wild-type NRAS^19–24^. A subsequent analysis of 1D free energy profiles revealed the barriers for the free energies dependent of *ϕ* and *d* are in the range of 8 to 9 kJ/mol. When GTP is hydrolyzed, accompanied by the dissociation of *γ*-phosphate, GDP-bound wild-type NRAS enters a so called called “GDP-specific conformation” state^4,62,65,66^, with the SW-I/II region largely opened. The free energy barrier to maintain this “GDP-specific conformation” state is approximately 22 kJ/mol, around 3 times larger than GTP-bound wild-type NRAS case. The NRAS Q61 mutation showed obvious differences in conformational changes compared with the wild type case. The most noticeable difference is related to the distance *d*, although the angles *ϕ* for the mutated species also tend to show smaller numbers in comparison with the wild-type case NRAS. For the positively charged mutations Q61R and Q61K, there exist large free energy barriers of 32 kJ/mol and 25 kJ/mol respectively, which corresponds to the strong HBs between positively charged R61 (K61) and negatively charged *γ*-phosphate. In a previous work^24^, we estimated by unbiased simulations the overall barriers of HBs between GDP-NRAS-R61(K61) and aqueous solutions to be of 8.9 *k*_*B*_*T* (23 kJ/mol) and 5.4 *k*_*B*_*T* (14 kJ/mol) respectively. This indicates that in the case of GDP-bound NRAS-Q61R(K)^24^, the SW-II can be separated from the protein surfaces, meanwhile the “GTP-bound” will maintain tight conformations in unbiased simulations.

In summary, benefiting from WTM simulations, we evaluated the FES of each NRAS case (rare GDP-bound NRAS-Q61 mutation cases are excluded) and accurately assessed the free energy barriers for the NRAS conformational changes and the effect of Q61 mutations on it. Meanwhile, through the WTM, we also calculated the free energy barriers for the interactions of R61(K61)-GTP, and L61 with the hydrophobic pocket located on *α*-helix3 (approximately 18 kJ/mol), which could not be accurately evaluated by the unbiased simulations. We can conclude that the methodology and results presented here can be useful tools to provide the detailed atomic-level structural information for the development/optimization of specific inhibitors for NRAS-Q61 mutations in the further investigation.

## Acknowledgements

We thank financial support provided by the Spanish Ministry of Science, Innovation and Universities (project number PGC2018-099277-B-C21, funds MCIU/AEI/FEDER, UE). This publication is a part of the I+D+i project Reference PID2021-124297NB-C32, founded by MCIN/ AEI/10.13039/501100011033/ and “FEDER Una manera de hacer Europa”. Zheyao Hu is a Ph.D. fellow from the Chinese Scholarship Council (grant 202006230070). J.M. thanks the *Generalitat de Catalunya* for the support through the grant *Grup de Recerca SGR-Cat2021 Condensed, Complex and Quantum Matter Group* reference 2021SGR-01411 and to the Polytechnic University of Catalonia-Barcelona Tech through the funding AGRUPS. Computational resources awarded by the Barcelona Supercomputing Center-Spanish Supercomputing Network (grant BCV-2024-2-0005 “Simulation and modelling of biosystems with applications to medicine (1): drug design for the treatment of cancers of the RAS family”) are also acknowledged.

## Notes and references

1 J. Cherfils and M. Zeghouf, Physiological reviews, 2013, 93, 269–309.

2 J. M. Ostrem and K. M. Shokat, Nature reviews Drug discovery, 2016, 15, 771–785.

3 W. Kolch, D. Berta and E. Rosta, Biochemical Journal, 2023, 480, 1–23.

4 I. R. Vetter and A. Wittinghofer, Science, 2001, 294, 1299– 1304.

5 N. Normanno, S. Tejpar, F. Morgillo, A. De Luca, E. Van Cutsem and F. Ciardiello, Nature reviews Clinical oncology, 2009, 6, 519–527.

6 C. M. Ardito, B. M. Grüner, K. K. Takeuchi, C. Lubeseder-Martellato, N. Teichmann, P. K. Mazur, K. E. DelGiorno, E. S. Carpenter, C. J. Halbrook, J. C. Hall et al., Cancer cell, 2012, 22, 304–317.

7 C. Navas, I. Hernández-Porras, A. J. Schuhmacher, M. Sibilia, C. Guerra and M. Barbacid, Cancer cell, 2012, 22, 318–330.

8 K. W. Wood, C. Sarnecki, T. M. Roberts and J. Blenis, Cell, 1992, 68, 1041–1050.

9 L. R. Howe, S. J. Leevers, N. Gómez, S. Nakielny, P. Cohen and C. J. Marshall, Cell, 1992, 71, 335–342.

10 P. Rodriguez-Viciana, P. H. Warne, R. Dhand, B. Vanhaesebroeck, I. Gout, M. J. Fry, M. D. Waterfield and J. Downward, Nature, 1994, 370, 527–532.

11 S. Li, A. Balmain and C. M. Counter, Nature Reviews Cancer, 2018, 18, 767–777.

12 A. R. Moore, S. C. Rosenberg, F. McCormick and S. Malek, Nat. Rev. Drug Discov., 2020, 19, 533–552.

13 I. A. Prior, F. E. Hood and J. L. Hartley, Cancer Res., 2020, 80, 2969–2974.

14 D. B. Johnson and I. Puzanov, Current treatment options in oncology, 2015, 16, 1–12.

15 T. Randic, I. Kozar, C. Margue, J. Utikal and S. Kreis, Cancer treatment reviews, 2021, 99, 102238.

16 D. Schadendorf, A. C. Van Akkooi, C. Berking, K. G. Griewank, R. Gutzmer, A. Hauschild, A. Stang, A. Roesch and S. Ugurel, The Lancet, 2018, 392, 971–984.

17 P. Nagarajan, M. M. Asgari, A. C. Green, S. M. Guhan, S. T. Arron, C. M. Proby, D. E. Rollison, C. A. Harwood and A. E. Toland, Clinical Cancer Research, 2019, 25, 2379–2391.

18 C. Posch and S. Ortiz-Urda, Oncotarget, 2013, 4, 494.

19 M. Frech, T. Darden, L. Pedersen, C. Foley, P. Charifson, M. Anderson and A. Wittinghofer, Biochemistry, 1994, 33, 3237–3244.

20 R. H. Tichauer, G. Favre, S. Cabantous, G. Landa, A. Hemeryck and M. Brut, Biophysical journal, 2018, 115, 1417–1430.

21 E. T. Novelli, J. T. First and L. J. Webb, Biochemistry, 2018, 57, 6356–6366.

22 R. H. Tichauer, G. Favre, S. Cabantous and M. Brut, The Journal of Physical Chemistry B, 2019, 123, 3935–3944.

23 K. A. Maegley, S. J. Admiraal and D. Herschlag, Proceedings of the National Academy of Sciences, 1996, 93, 8160–8166.

24 Z. Hu and J. Martí, Computational and structural biotechnology journal, 2024.

25 I. V. Fedorenko, G. T. Gibney and K. S. Smalley, Oncogene, 2013, 32, 3009–3018.

26 D. B. Johnson, C. M. Lovly, M. Flavin, K. S. Panageas, G. D. Ayers, Z. Zhao, W. T. Iams, M. Colgan, S. DeNoble, C. R. Terry et al., Cancer immunology research, 2015, 3, 288–295.

27 L. E. Davis, S. C. Shalin and A. J. Tackett, Cancer biology & therapy, 2019, 20, 1366–1379.

28 D. Liu, Y. Mao, X. Gu, Y. Zhou and D. Long, Proceedings of the National Academy of Sciences, 2021, 118, e2024725118.

29 A. Barducci, M. Bonomi and M. Parrinello, Wiley Interdisciplinary Reviews: Computational Molecular Science, 2011, 1, 826–843.

30 G. Bussi and A. Laio, Nature Reviews Physics, 2020, 2, 200– 212.

31 J. McCarty and M. Parrinello, The Journal of chemical physics, 2017, 147, year.

32 W. L. Jorgensen, J. Chandrasekhar, J. D. Madura, R. W. Impey and M. L. Klein, The Journal of chemical physics, 1983, 79, 926–935.

33 S. Jo, T. Kim, V. G. Iyer and W. Im, Journal of Computational Chemistry, 2008, 29, 1859–1865.

34 B. R. Brooks, C. L. Brooks III, A. D. Mackerell Jr, L. Nilsson, R. J. Petrella, B. Roux, Y. Won, G. Archontis, C. Bartels, S. Boresch et al., Journal of computational chemistry, 2009, 30, 1545–1614.

35 J. Lee, X. Cheng, J. M. Swails, M. S. Yeom, P. K. Eastman, J. A. Lemkul, S. Wei, J. Buckner, J. C. Jeong and Y. e. a. Qi, Journal of chemical theory and computation, 2016, 12, 405–413.

36 J. Huang and A. D. MacKerell Jr, Journal of computational chemistry, 2013, 34, 2135–2145.

37 A. Kouranov, L. Xie, J. de la Cruz, L. Chen, J. Westbrook, P. E. Bourne and H. M. Berman, Nucleic acids research, 2006, 34, D302–D305.

38 H. J. Berendsen, D. van der Spoel and R. van Drunen, Computer Physics Communications, 1995, 91, 43–56.

39 H. M. Senn and W. Thiel, in Atomistic approaches in modern biology, Springer, 2006, pp. 173–290.

40 P. L. Geissler, C. Dellago, D. Chandler, J. Hutter and M. Parrinello, Science, 2001, 291, 2121–2124.

41 C. Dellago, P. G. Bolhuis and P. L. Geissler, Advances in chemical physics, 2002, 123, 1–78.

42 J. Henin, G. Fiorin, C. Chipot and M. L. Klein, Journal of Chemical Theory and Computation, 2009, 6, 35–47.

43 C. Bartels and M. Karplus, Journal of Computational Chemistry, 1997, 18, 1450–1462.

44 G. Bussi, F. L. Gervasio, A. Laio and M. Parrinello, Journal of the American Chemical Society, 2006, 128, 13435–13441.

45 M. Deighan, M. Bonomi and J. Pfaendtner, Journal of Chemical Theory and Computation, 2012, 8, 2189–2192.

46 J. C. Palmer, R. Car and P. G. Debenedetti, Faraday Discussions, 2013, 167, 77–94.

47 S. Haldar, P. Kührová, P. Banáš, V. Spiwok, J. Sponer, P. Hobza and M. Otyepka, Journal of Chemical Theory and Computation, 2015, 11, 3866–3877.

48 J. Martí, Molecular Simulation, 2018, 44, 1136–1146.

49 H. Lu and J. Marti, Scientific reports, 2020, 10, 9235.

50 H. Lu and J. Marti, The Journal of Physical Chemistry Letters, 2020, 11, 9938–9945.

51 J. Martí and H. Lu, International journal of molecular sciences, 2021, 22, 2842.

52 Z. Hu and J. Marti, Molecular Physics, 2024, e2316883.

53 M. Bonomi, D. Branduardi, G. Bussi, C. Camilloni, D. Provasi, P. Raiteri, D. Donadio, F. Marinelli, F. Pietrucci and R. A. e. a. Broglia, Computer Physics Communications, 2009, 180, 1961–1972.

54 G. A. Tribello, M. Bonomi, D. Branduardi, C. Camilloni and G. Bussi, Computer Physics Communications, 2014, 185, 604– 613.

55 G. Bussi, D. Branduardi et al., Rev. Comput. Chem, 2015, 28, 1–49.

56 A. Barducci, G. Bussi and M. Parrinello, Physical review letters, 2008, 100, 020603.

57 P. Hošek and V. Spiwok, Computer Physics Communications, 2016, 198, 222–229.

58 W. Humphrey, A. Dalke and K. Schulten, Journal of molecular graphics, 1996, 14, 33–38.

59 G. Bussi, A. Laio and M. Parrinello, Physical review letters, 2006, 96, 090601.

60 M. Bonomi, C. Camilloni and M. Vendruscolo, Scientific reports, 2016, 6, 31232.

61 H. Lu and J. Marti, Membranes, 2020, 10, 364.

62 Z. Hu and J. Marti, International journal of molecular sciences, 2022, 23, 13865.

63 R. Nussinov, C.-J. Tsai and H. Jang, Cancer research, 2018, 78, 593–602.

64 A. R. Moore, S. C. Rosenberg, F. McCormick and S. Malek, Nat. Rev. Drug Discov.

65 P. Grudzien, H. Jang, N. Leschinsky, R. Nussinov and V. Gaponenko, Journal of Molecular Biology, 2022, 434, 167695.

66 S. Lu, H. Jang, S. Muratcioglu, A. Gursoy, O. Keskin, R. Nussinov and J. Zhang, Chemical reviews, 2016, 116, 6607–6665.

67 J. P. McGrath, D. J. Capon, D. V. Goeddel and A. D. Levinson, Nature, 1984, 310, 644–649.

68 R. W. Sweet, S. Yokoyama, T. Kamata, J. R. Feramisco, M. Rosenberg and M. Gross, Nature, 1984, 311, 273–275.

69 M. Trahey and F. McCormick, Science, 1987, 238, 542–545.

70 H. Adari, D. R. Lowy, B. M. Willumsen, C.J. Der and F. McCormick, Science, 1988, 240, 518–521.

71 J. P. Jambeck and A. P. Lyubartsev, The Journal of Physical Chemistry Letters, 2013, 4, 1781–1787.

72 B. Roux, Biophysical journal, 1999, 77, 139–153.

