## Supporting information for "Quantitative comparison of the structural differences between NRAS and its mutations by well-tempered metadynamics simulations"

Zheyao Hu[a] and Jordi Martí*[a]

### 1 The characteristics of systems and convergence of well-tempered metadynamics simulations

The five classes of NRAS proteins that have been considered in this work such as GTP-bound wild-type NRAS, NRAS-Q61R, NRAS-Q61K, NRAS-Q61L and GDP-bound wild-type NRAS. The initial setups of these systems are same with the previous work[1]. The initial structure of NRAS-WT were obtained from Protein Data Bank (structures 6zio.pdb and 5uhv.pdb, respectively). The energy minimization and equilibration of the simulation systems have been reached under the same conditions as before[1]. Afterwards, five long well-tempered metadynamics simulations of 3 $\mu s$ each have been simulated in order to explore the free-energy surface of NRAS and its mutants.

Concerning the reliability of the simulations reported in the present work, we can use several criteria. Three of the most efficient and reliable are: (1) the time evolution of the fluctuations of the hills of the biased potential along the full trajectory; (2) the time cumulative average of 1D free energy profiles as defined in main text (Section 3.4) and (3) considering the so-called block analysis of the average error along free-energy profiles as a function of the block length. Here, for convenience, we mainly report the time evolution of the fluctuations of the hills (Fig. 1) and the block analysis of the average error along free-energy profiles as a function of the block length (Fig. 2).

The evolution of the hills is presented in Fig. 1. We observe an overall decreasing behaviour, eventually including low-height spikes. In all cases, the height of the biased potential decreased accordingly along the simulation runs. In all cases, a quasi-flat profile is already reached. This is a clear indication of the convergence of the WTM runs. Moreover, from the results of Fig. 2, where we report the size of the average error in the free-energy profiles calculated from different sets of well-tempered metadynamics simulations as a function of the block size. The full length of the metadynamics trajectories were taken into account. As expected, the errors increase with the block length until they reach a plateau in all cases. For this calculation, scripts provided by the PLUMED project[2,3] have been employed.

**Table 1** The data of 2D free-energy landscapes minima: GTP-bound wild-type NRAS

| Basins | $d$ [nm] | $\phi$ [rad] | free energy [kJ/mol] |
|---|---|---|---|
| A | 0.7 | 2.1 | 0.0 |
| B | 0.7 | 1.7 | 1.4 |
| C | 0.9 | 2.1 | 1.6 |
| D | 0.7 | 0.5 | 2.4 |
| E | 0.5 | 1.8 | 2.7 |
| F | 0.9 | 0.5 | 3.0 |
| G | 1.2 | 1.7 | 6.3 |
| H | 1.7 | 0.7 | 11.6 |
| I | 0.5 | 0.7 | 13.6 |

**Table 2** The data of 2D free-energy landscapes minima: GDP-bound wild-type NRAS

| Basins | $d$ [nm] | $\phi$ [rad] | free energy [kJ/mol] |
|---|---|---|---|
| A | 1.2 | 2.3 | 0.0 |
| B | 0.7 | 2.3 | 0.0 |
| C | 0.5 | 2.3 | 12.5 |
| D | 1.0 | 0.4 | 14.2 |
| E | 0.7 | 0.5 | 15.6 |
| G | 0.5 | 0.4 | 18.6 |

### 2 The data of 2D free-energy landscapes minima

The detailed coordinates of minima and the corresponding free energy values are reported in Table 1 to Table 5. In order to directly compute the height of free-energy barriers, the absolute minima are set equal to zero.

a‡Department of Physics, Technical University of Catalonia-Barcelona Tech, B5-209 Northern Campus, Jordi Girona 1-3, 08034 Barcelona, Catalonia, Spain.

a‡Department of Physics, Technical University of Catalonia-Barcelona Tech, B5-209 Northern Campus, Jordi Girona 1-3, 08034 Barcelona, Catalonia, Spain.

*Corresponding author

‡These authors contributed equally to this work

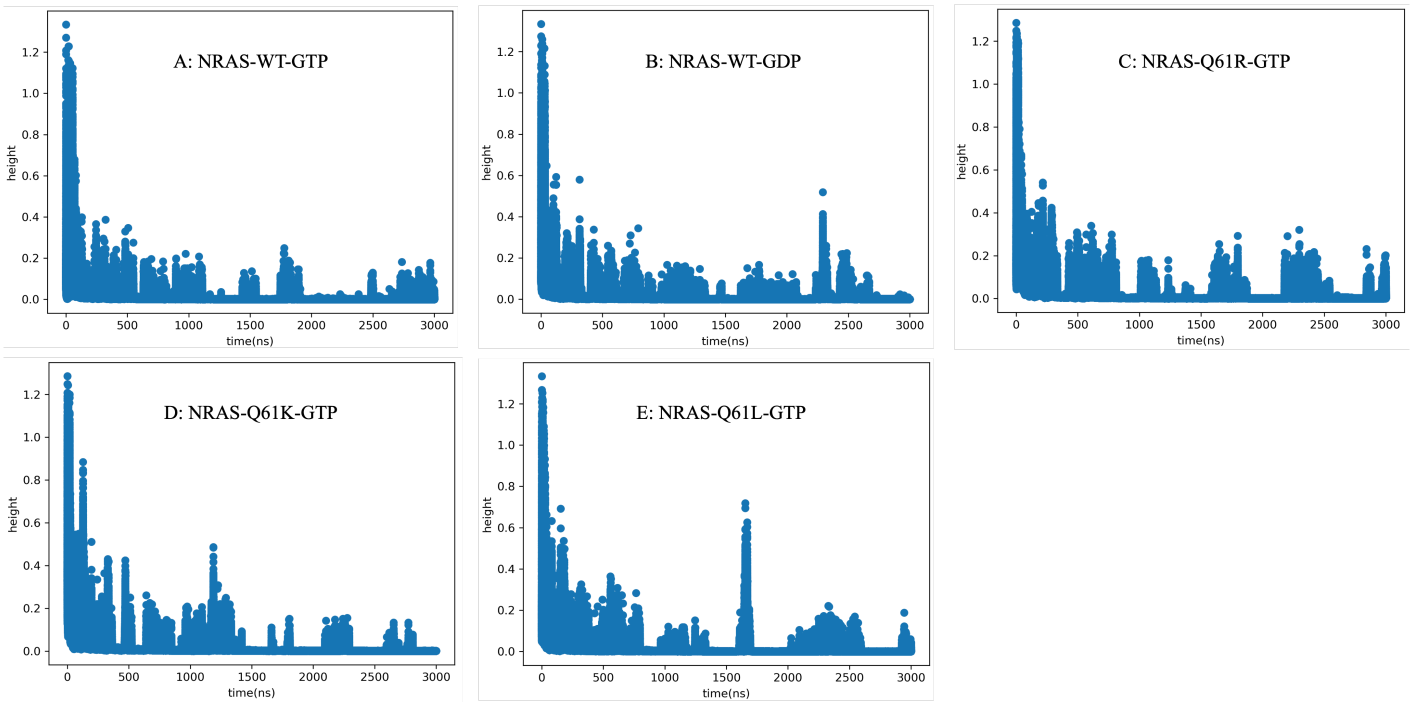


**Fig. 1** Height of the hills of the Gaussian-biased potential as a function of time. All profiles correspond to the WTM simulations of each NRAS systems.

**Table 3** The data of 2D free-energy landscapes minima: GTP-bound NRAS-Q61R

| Basins | $d$ [nm] | $\phi$ [rad] | free energy [kJ/mol] |
|---|---|---|---|
| A | 0.4 | 1.1 | 0.0 |
| B | 0.4 | 0.4 | 2.7 |
| C | 1.4 | 0.5 | 3.1 |
| D | 0.4 | 1.6 | 5.4 |

**Table 4** The data of 2D free-energy landscapes minima: GTP-bound NRAS-Q61K

| Basins | $d$ [nm] | $\phi$ [rad] | free energy [kJ/mol] |
|---|---|---|---|
| A | 0.4 | 0.5 | 0.0 |
| B | 1.4 | 0.7 | 3.5 |
| C | 1.4 | 2.1 | 7.7 |
| D | 0.4 | 2.2 | 8.0 |
| E | 0.4 | 0.9 | 8.4 |
| F | 0.4 | 1.3 | 9.4 |
| G | 0.6 | 0.8 | 22.6 |

**Table 5** The data of 2D free-energy landscapes minima: GTP-bound NRAS-Q61L

| Basins | $d$ [nm] | $\phi$ [rad] | free energy [kJ/mol] |
|---|---|---|---|
| A | 0.3 | 0.8 | 0.0 |
| B | 1.5 | 1.1 | 9.1 |
| C | 1.0 | 1.1 | 12.9 |

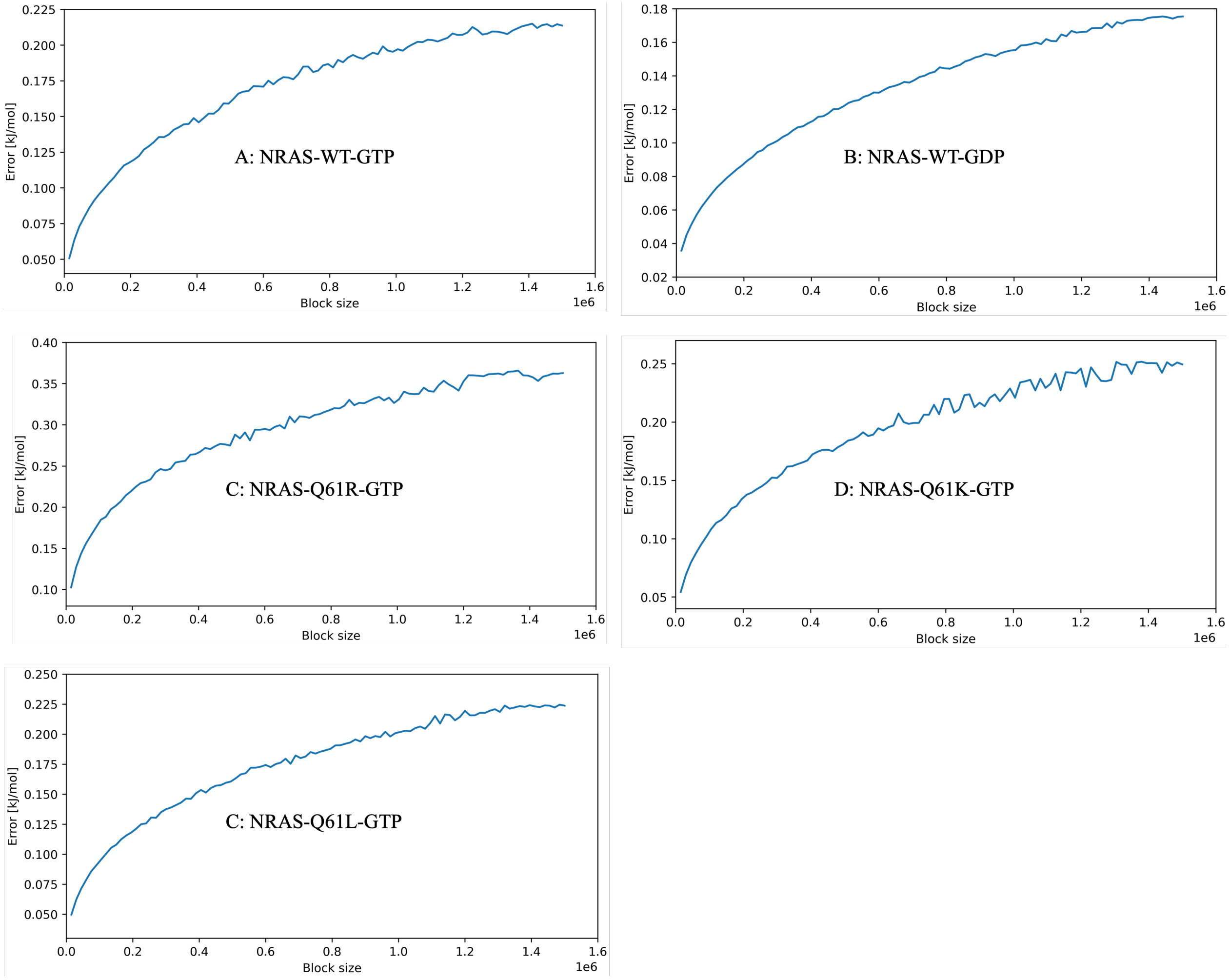


**Fig. 2** Block analysis of the average errors along the free-energy profiles for CVs as a function of the block length. (A) NRAS-WT-GTP case: angle $\phi$; (B) NRAS-WT-GDP case: angle $\phi$; (C) NRAS-Q61R-GTP case: angle $\phi$; (D) NRAS-Q61K-GTP case: distance $d$; and (E) NRAS-Q61L-GTP case: angle $\phi$. The entire metadynamics simulation (3.0 $\mu$s for each system) is taken into account.

### Notes and references


1 Z. Hu and J. Martí, *Computational and structural biotechnology journal*, 2024.

2 M. Bonomi, D. Branduardi, G. Bussi, C. Camilloni, D. Provasi, P. Raiteri, D. Donadio, F. Marinelli, F. Pietrucci and R. A. e. a. Broglia, *Computer Physics Communications*, 2009, **180**, 1961–1972.

3 G. A. Tribello, M. Bonomi, D. Branduardi, C. Camilloni and G. Bussi, *Computer Physics Communications*, 2014, **185**, 604–613.
